# Evidence that a major subpopulation of fall armyworm found in the Western Hemisphere is rare or absent in Africa, which may limit the range of crops at risk of infestation

**DOI:** 10.1101/482935

**Authors:** Rodney N. Nagoshi

**Affiliations:** Center for Medical, Agricultural and Veterinary Entomology, United States Department of Agriculture, Agricultural Research Service, Gainesville, Florida, United States of America.

## Abstract

The introduction and establishment of fall armyworm (*Spodoptera frugiperda*) in Africa presents a major threat to agriculture in that continent and potentially to the entire Eastern Hemisphere. The species is subdivided into two subpopulations called the R-strain and C-strain that differ in their distribution on different plant hosts. This means that the scope of the economic risk posed by invasive fall armyworm is influenced by whether one or both strains are present. Multiple studies have found mitochondrial markers diagnostic of the two strains throughout Africa but there is substantial disagreement with a nuclear strain marker that makes conclusions about strain composition uncertain. In this study the issue of whether both strains are present in Africa was tested by an assay that can detect strain-biased mating behaviors. Western Hemisphere fall armyworm consistently showed evidence of strain-specific assortative mating in the field that was not found in surveys from multiple locations in Africa. The absence of strain mating biases and the disagreements between the strain diagnostic genetic markers indicates that the R-strain is rare (<1% of the population) or absent in Africa. Instead, it appears that the African fall armyworm populations are dominated by two groups, the C-strain and the descendants of interstrain hybrids. These results suggest that plant hosts associated with the R-strain may not be at high risk of fall armyworm infestation in Africa.

## Introduction

Fall armyworm (*Spodoptera frugiperda*) is an important agricultural pest of the Western Hemisphere that has recently spread into Africa where it has the potential for significant economic impact [1]. The species has a broad host range having been found on greater than 80 plant species and cultivars [2]. However, infestations of agricultural importance are limited to a few crops in the Western Hemisphere, indicating that while capable of being broadly polyphagous, fall armyworm tends to be more specific in its host usage. The species can be subdivided into two subpopulations defined by their differential distribution in the field [3]. The C-strain (historically designated the corn-strain) is typically found on corn, sorghum, and cotton, while the R-strain (rice-strain) predominates on rice, alfalfa, pasture grasses, and millet, though subsequent observations indicate that the preference to rice may be variable [4-7]. There is genetic evidence that both strains are present in Africa suggesting that a variety of crop systems are at risk [8-11]. However, reports of fall armyworm damage to date have been primarily limited to C-strain preferred hosts, specifically corn and sorghum [12]. There are some anecdotal reports of infestations in R-strain host plants, but whether these occur consistently or incur significant economic damage has yet to be documented.

The fall armyworm strains were originally defined by the differential distribution of genetic markers among host plants in larval field collections [3, 13]. Most commonly used for population studies are mitochondrial haplotypes, with those defined by polymorphisms in the *Cytochrome oxidase subunit I* gene (*COI*) the best characterized [14-16]. Strain markers from nuclear DNA have been difficult to isolate and are currently limited to a small number of polymorphic loci located on the *Z* sex chromosome [17-19]. One such set of *Z*-linked markers are found in the *Triosephosphate isomerase* gene (*Tpi*), which is thought to have a general housekeeping function [20] and is highly conserved in noctuid moths [18]. The *Tpi* gene may be linked to loci important for speciation as polymorphisms can distinguish between the sibling species *Helicoverpa armigera* (Hübner) and *H. zea* (Boddie) and two "races" of *Ostrinia nubialis* (Hübner) [21, 22]. In fall armyworm the correspondence between the *Tpi* strain markers and host plants tends to be higher than that observed with mitochondrial markers, suggesting that *Tpi* is the more accurate indicator of strain identity [18, 23].

The preferential association of C-strain (*COI*-CS and *Tpi*C) and R-strain (*COI*-RS and *Tpi*R) markers to larvae found on different host plants has been demonstrated in both Americas, indicating a hemispheric distribution of the two strains [5, 6]. However, the correspondence between marker and plant host is not absolute. Inconsistencies include instances where specimens collected from a particular plant host do not have the expected strain marker [4, 18, 24, 25], as well as disagreements between the mitochondrial and nuclear markers [13, 18, 26]. Such inconsistencies could result from interstrain mating, where the resulting hybrids and their derivatives would have variable marker configurations and unknown plant host preferences.

Field studies of the two strains are difficult because of their morphological and behavioral similarities. Strain-specific differences have been reported for mating behavior [27-31], pesticide resistance [32-34], and pheromone composition [35-37]. However, fall armyworm can exhibit considerable phenotypic variation between regional populations and laboratory colonies that are independent of strain differences. For example, differential response to pheromone lures was found to vary by geography as much as by strain [38] and the observation of strain-specific mating behaviors can be colony dependent [39]. There are reports that South American fall armyworm display significant strain-differences in wing shape and size [40, 41], but it is not yet known whether this phenotypic distinction applies to populations from other locations. At this time the strains are considered to be morphologically indistinguishable with genetic markers remaining the most reliable means of strain identification.

Pheromone trap collections typically contain both host strains indicating a sympatric distribution. This would seem to require the existence of reproductive barriers between strains to maintain their continued divergence. Several mechanisms have been reported based primarily on laboratory studies. These include strain-specific assortative mating [28, 29] attributed to differences in female pheromone composition [36, 42] and the timing of mating behavior during scotophase [28, 30, 31]. Post-mating barriers have also been observed in the form of lower viability or fertility of interstrain hybrids [29, 39]. However, at this time there has been no conclusive demonstration that any of these mechanisms are actually restricting interstrain hybridization in the field.

This uncertainty is in large part due to the difficulty in identifying interstrain hybrids given the morphological equivalence of the two parental strains. For this reason, we recently developed a strategy for estimating the frequency of interstrain hybrid formation in field populations that used neutral single-base polymorphisms (SNPs) within the *Tpi* coding region that differ in their degree of strain-specificity [43]. In principle, the frequency of heterozygosity at the sites with high strain-specificity should be indicative of interstrain hybridization, while heterozygosity at sites with low specificity will reflect successful mating both within and between strains. Therefore, comparisons between the two should provide a direct measure of the frequency of interstrain relative to intrastrain hybridization. The results obtained from collections at multiple sites in the United States were consistent with at least a 4-fold reduction in hybridization frequency between strains versus within strains [43].

In this paper, the distribution of genetic markers in Africa was examined in more detail and compared with observations from the Western Hemisphere. Previously reported disagreements between the *COI* and *Tpi* markers in African populations [9, 10] were quantified and reinterpreted from the perspective of interstrain hybrids. The presence of the R-strain and the possible impact of hybrids on mating behavior were examined by the application of the *Tpi* heterozygosity methodology on the African fall armyworm populations. We first tested whether this methodology was sufficiently robust to detect strain-specific heterozygosity differences in larval collections where gender is not known, as well as with databases where multiple collections are pooled. The analysis was then applied to pheromone trap collections and larval samplings from Africa. The implications of the results to the strain composition of the African fall armyworm population are discussed.

## Materials and Methods

Fall armyworm were collected from pheromone traps (adult males) and larvae from corn or sorghum host plants (gender unknown) at various sites in the Western Hemisphere and Africa (Fig. 1, Table 1). Collections have been previously described, with their archived DNA preparations or additional specimens used for the new analysis in this study. All specimens were stored at −20°C until DNA preparation.

**Figure 1.**
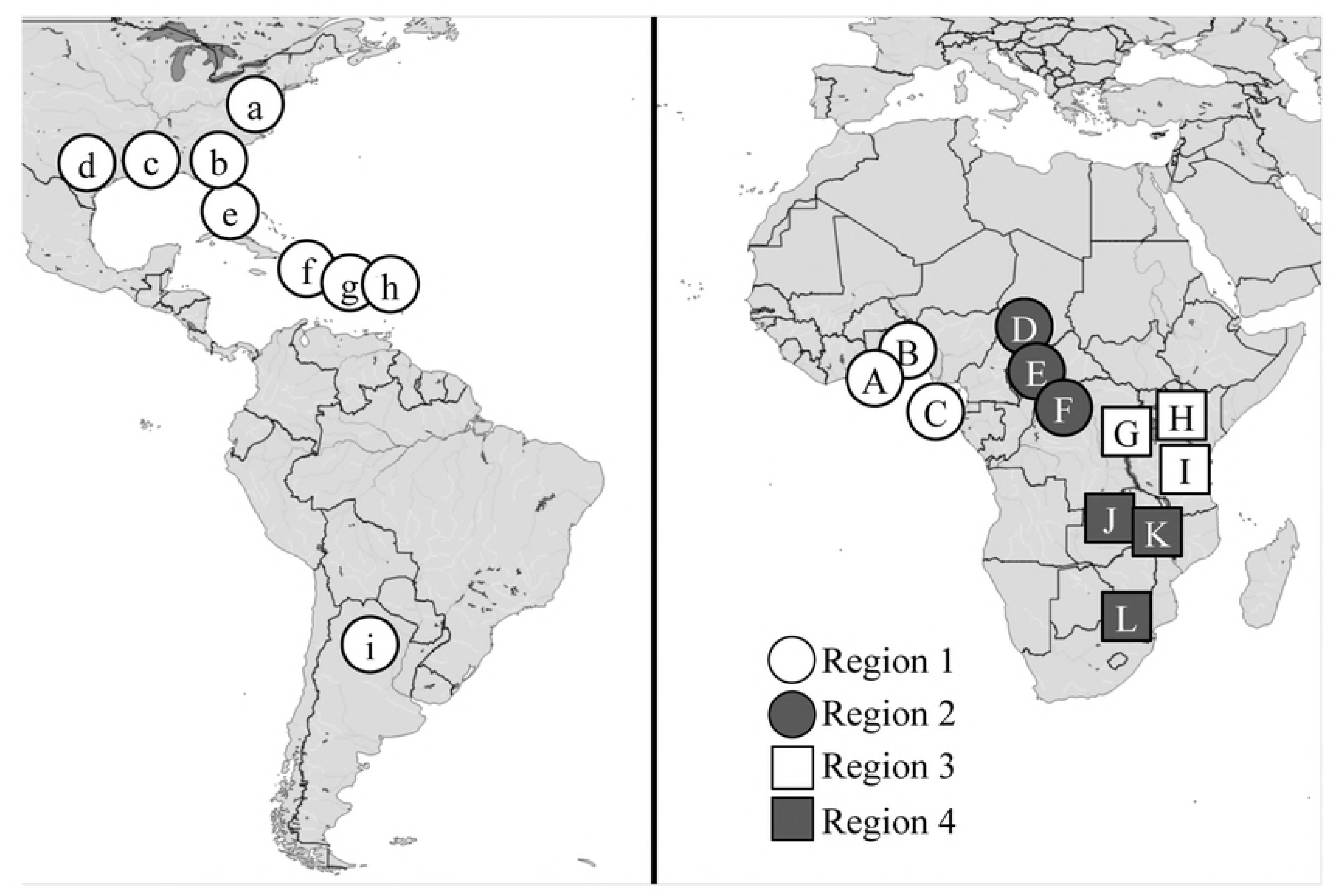
Map of the Western Hemisphere and Africa showing locations of fall armyworm collections. Letters refer to Table 1 (parenthesis). Africa collections were grouped into regions designated by shapes and shadings.

**Table 1.**
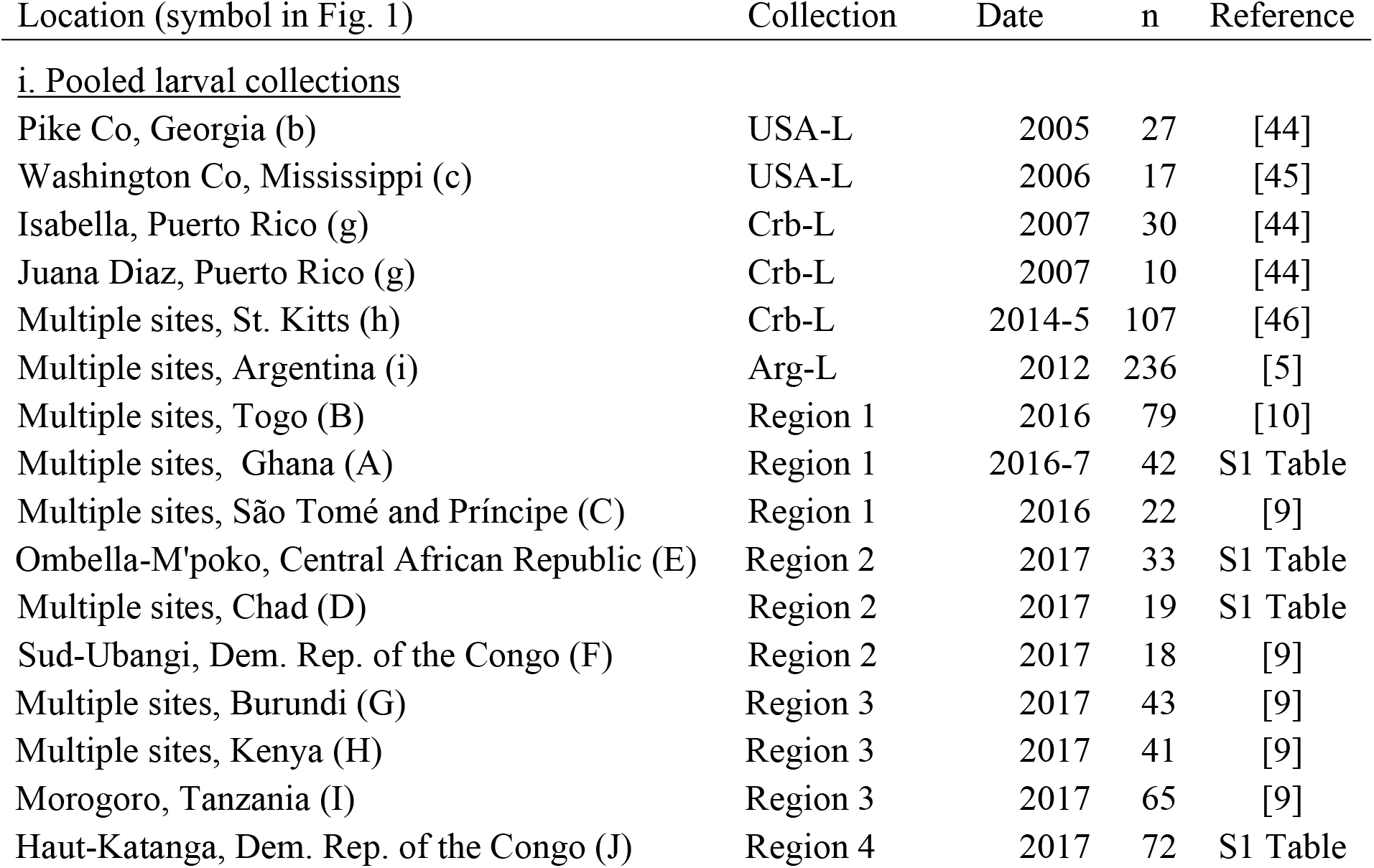
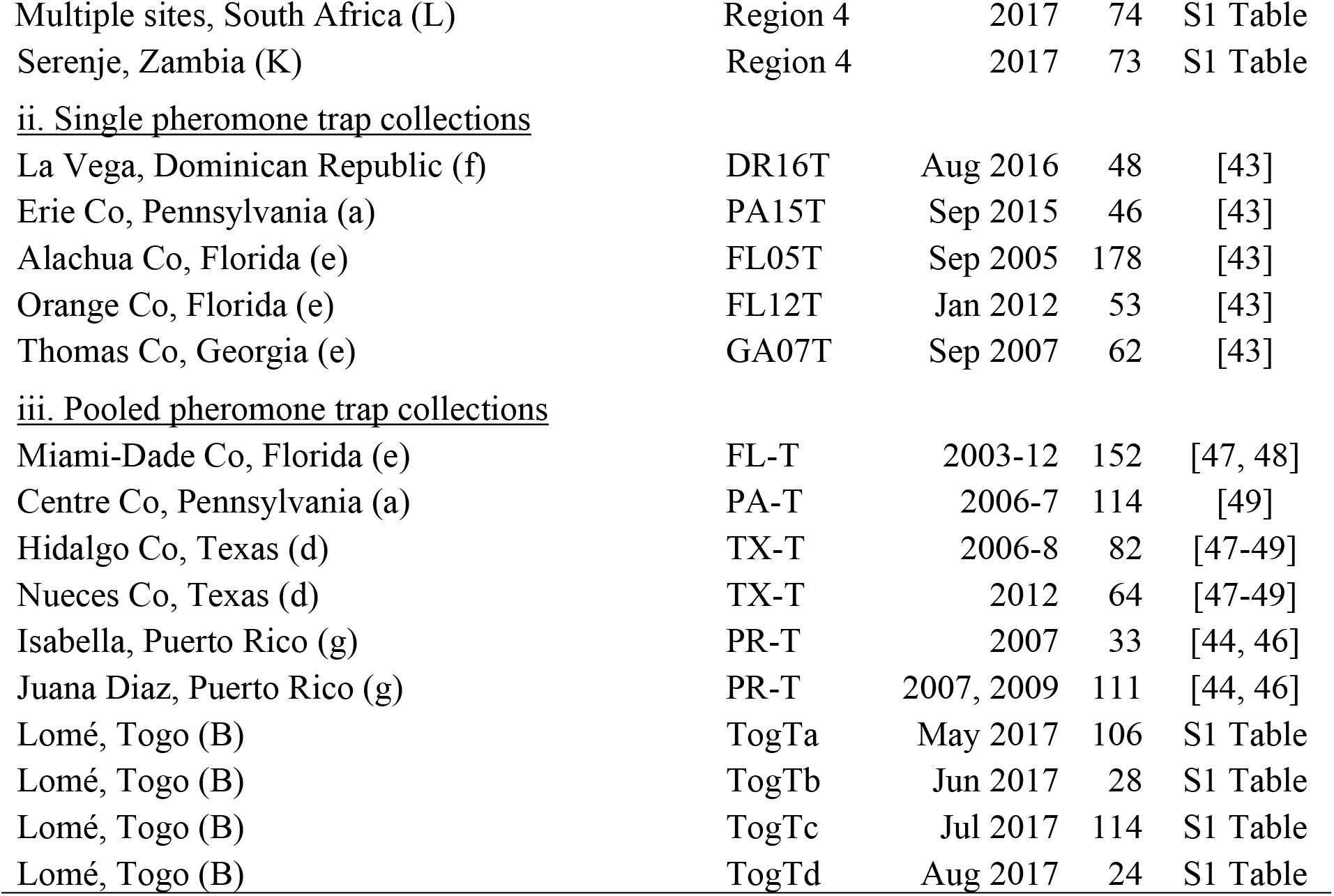
Fall armyworm collections.

| Location (symbol in Fig. 1) | Collection | Date | n | Reference |
| --- | --- | --- | --- | --- |
| <u>i. Pooled larval collections</u> |  |  |  |  |
| Pike Co, Georgia (b) | USA-L | 2005 | 27 | [44] |
| Washington Co, Mississippi (c) | USA-L | 2006 | 17 | [45] |
| Isabella, Puerto Rico (g) | Crb-L | 2007 | 30 | [44] |
| Juana Diaz, Puerto Rico (g) | Crb-L | 2007 | 10 | [44] |
| Multiple sites, St. Kitts (h) | Crb-L | 2014-5 | 107 | [46] |
| Multiple sites, Argentina (i) | Arg-L | 2012 | 236 | [5] |
| Multiple sites, Togo (B) | Region 1 | 2016 | 79 | [10] |
| Multiple sites, Ghana (A) | Region 1 | 2016-7 | 42 | S1 Table |
| Multiple sites, São Tomé and Príncipe (C) | Region 1 | 2016 | 22 | [9] |
| Ombella-M'poko, Central African Republic (E) | Region 2 | 2017 | 33 | S1 Table |
| Multiple sites, Chad (D) | Region 2 | 2017 | 19 | S1 Table |
| Sud-Ubangi, Dem. Rep. of the Congo (F) | Region 2 | 2017 | 18 | [9] |
| Multiple sites, Burundi (G) | Region 3 | 2017 | 43 | [9] |
| Multiple sites, Kenya (H) | Region 3 | 2017 | 41 | [9] |
| Morogoro, Tanzania (I) | Region 3 | 2017 | 65 | [9] |
| Haut-Katanga, Dem. Rep. of the Congo (J) | Region 4 | 2017 | 72 | S1 Table |
| Multiple sites, South Africa (L) | Region 4 | 2017 | 74 | S1 Table |
| Serenje, Zambia (K) | Region 4 | 2017 | 73 | S1 Table |
| ii. Single pheromone trap collections |  |  |  |  |
| La Vega, Dominican Republic (f) | DR16T | Aug 2016 | 48 | [43] |
| Erie Co, Pennsylvania (a) | PA15T | Sep 2015 | 46 | [43] |
| Alachua Co, Florida (e) | FL05T | Sep 2005 | 178 | [43] |
| Orange Co, Florida (e) | FL12T | Jan 2012 | 53 | [43] |
| Thomas Co, Georgia (e) | GA07T | Sep 2007 | 62 | [43] |
| iii. Pooled pheromone trap collections |  |  |  |  |
| Miami-Dade Co, Florida (e) | FL-T | 2003-12 | 152 | [47, 48] |
| Centre Co, Pennsylvania (a) | PA-T | 2006-7 | 114 | [49] |
| Hidalgo Co, Texas (d) | TX-T | 2006-8 | 82 | [47-49] |
| Nueces Co, Texas (d) | TX-T | 2012 | 64 | [47-49] |
| Isabella, Puerto Rico (g) | PR-T | 2007 | 33 | [44, 46] |
| Juana Diaz, Puerto Rico (g) | PR-T | 2007, 2009 | 111 | [44, 46] |
| Lomé, Togo (B) | TogTa | May 2017 | 106 | S1 Table |
| Lomé, Togo (B) | TogTb | Jun 2017 | 28 | S1 Table |
| Lomé, Togo (B) | TogTc | Jul 2017 | 114 | S1 Table |
| Lomé, Togo (B) | TogTd | Aug 2017 | 24 | S1 Table |

### DNA Preparation and PCR amplification

Individual specimens were homogenized in 1.5 ml of phosphate buffered saline (PBS, 20 mM sodium phosphate, 150 mM NaCl, pH 8.0) using a tissue homogenizer (PRO Scientific Inc., Oxford, CT) or hand-held Dounce homogenizer and the homogenate transferred to a 2-ml microcentrifuge tube. Cells and tissue were pelleted by centrifugation at 6000g for 5 min. at room temperature. The pellet was resuspended in 800 µl Genomic Lysis buffer (Zymo Research, Orange, CA) by vortexing and incubated at 55°C for 5 min. Debris was removed by centrifugation at 10,000 rpm for 3 min. The supernatant was transferred to a Zymo-Spin III column (Zymo Research, Orange, CA) and processed according to manufacturer’s instructions. The DNA preparation was increased to a final volume of 100 µl with distilled water. Genomic DNA preparations of fall armyworm samples from previous studies were stored at −20°C.

Polymerase chain reaction (PCR) amplification for each gene segment was done separately, using a 30-µl reaction mix containing 3 µl 10X manufacturer’s reaction buffer, 1 µl 10mM dNTP, 0.5 µl 20-µM primer mix, 1 µl DNA template (between 0.05-0.5 µg), 0.5 units Taq DNA polymerase (New England Biolabs, Beverly, MA) and water. The thermocycling program was 94°C (1 min), followed by 33 cycles of 92°C (30 s), 56°C (45 s), 72°C (45 s), and a final segment of 72°C for 3 min. Typically 96 PCR amplifications were performed at the same time using either 0.2-ml tube strips or 96 well microtiter plates. Primers were synthesized by Integrated DNA Technologies (Coralville, IA). Amplification of *CO1* used the primer pair *CO1-891F* (5’-TACACGAGCATATTTTACATC-3’) and *CO1-1472R* (5’-GCTGGTGGTAAATTTTGATATC-3’) to produce a 603-bp fragment. Amplification of the *Tpi* region was done with the primers *Tpi412F* (5’-CCGGACTGAAGGTTATCGCTTG-3’) and *Tpi1140R* (5’-GCGGAAGCATTCGCTGACAACC-3’) that spans a variable length intron to produce a fragment with a mean length of 500 bp.

For fragment isolations, 6 µl of 6X gel loading buffer was added to each amplification reaction and the entire sample run on a 1.8% agarose horizontal gel containing GelRed (Biotium, Hayward, CA) in 0.5X Tris-borate buffer (TBE, 45 mM Tris base, 45 mM boric acid, 1 mM EDTA pH 8.0). Fragments were visualized on a blue LED illuminator (Biotium, Hayward, CA) and cut out from the gel. Fragment isolation was performed using Zymo-Spin I columns (Zymo Research, Orange, CA) according to manufacturer’s instructions. DNA sequencing was performed directly from the gel purified PCR fragments by Sanger sequencing (Genewiz, South Plainfield, NJ).

### Determination of strain-identity using *COI*

Preliminary identification of strain identity involved *Eco*RV restriction enzyme (New England Biolabs, Beverly, MA) of the PCR product produced by the *CO1*-891F/1472R primers. A strain-specific polymorphism at site 1182 produces in an EcoRV recognition site in the COI-RS haplotype but not with COI-CS [6, 14, 25]. Uncut fragments were further analyzed by DNA sequencing to confirm C-strain identity.

### Analysis of *Tpi* polymorphisms

The single nucleotide polymorphisms (SNPs) used in this study are located in the fourth exon of the *Tpi* coding region. Three strain-specific SNPs are present, of which e4_183_ is representative and diagnostic in this study of the *Tpi*C and *Tpi*R haplotypes ([18], Fig. 2A). Two other SNPs are found that also show high variability but display much less strain-specificity than e4_183_ [43]. For the purposes of this study they were considered non-specific, with site e4_192_ serving as the representative for both. The polymorphism associated with all five SNPs is a choice between C or T, neither of which alters the presumptive amino acid sequence.

**Figure 2.**
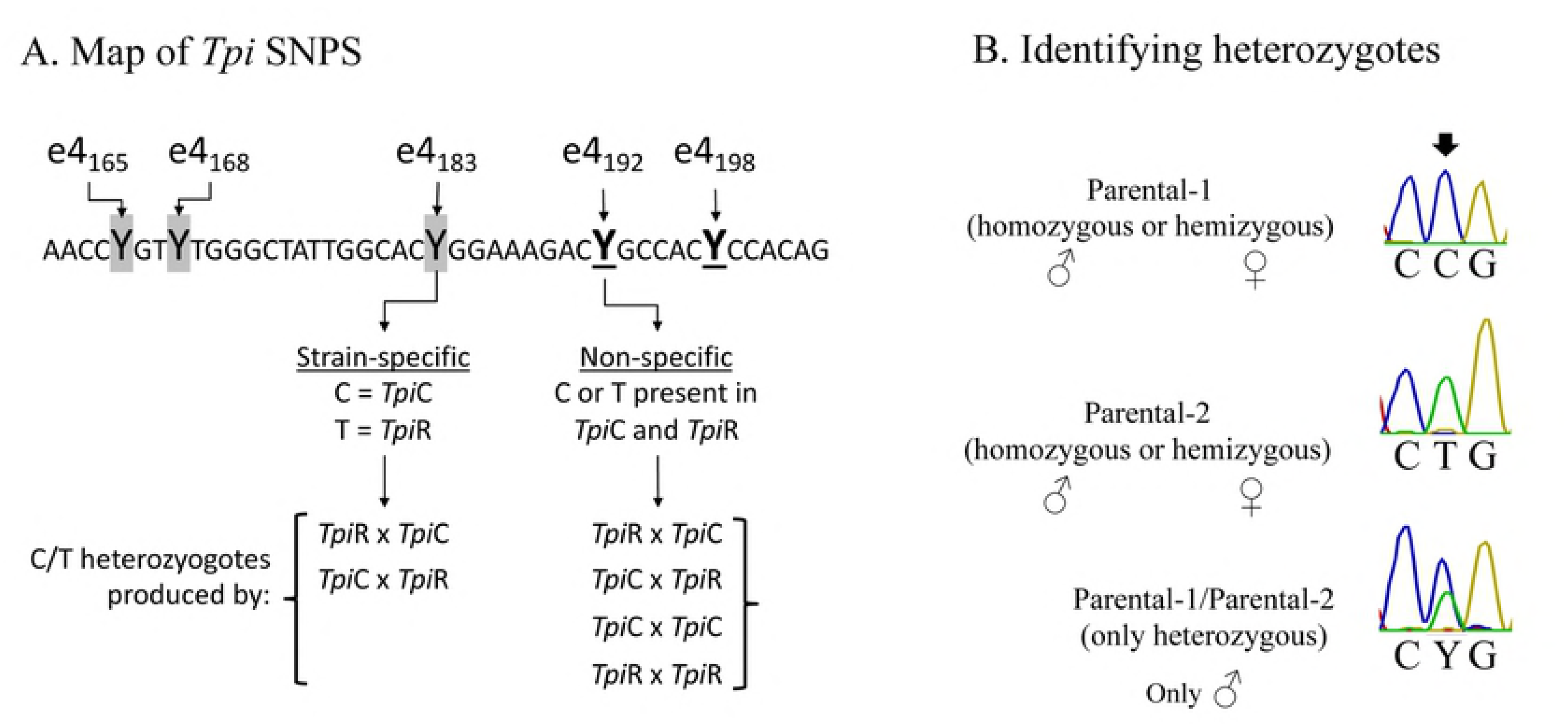
Map of selected *Tpi* SNPs that differ in strain-specificity including a list of the crosses that will produce heterozygotes at these sites and a description of the method to identify heterozygotes. (A) DNA sequence of the segment of the fourth exon of the coding region containing three strain-specific sites (shaded) and two non-specific sites (underlined and bold). All SNPs are associated with a C or T (designated Y by IUPAC convention). Below are the types of crosses that can produce heterozygotes at e4_183_ and e4_192_. (B) Example of DNA sequencing chromatographs diagnostic of the two parental alleles as present as a hemizygote or homozygote, and the chromatograph representing the heterozygous combination that shows an overlap of the parental curves.

The PCR amplified fragment produced by the primer pair *Tpi*412F/1140R contains both SNP sites and can be simultaneously read from a single sequencing run using *Tpi*412F as primer. The *Tpi* gene is *Z*-linked and so is hemizygous in females (*ZW*) and either homozygous or heterozygous in males (*ZZ*) with respect to the e4_183_ and e4_192_ SNPs. Sequencing of the PCR product amplified from genomic DNA can distinguish between homozygotes and heterozygotes, with the latter identified by overlapping chromatographs at the polymorphic site (Fig. 2B). However, this method cannot distinguish hemizygotes from homozygotes, which is not relevant for trap collections where all specimens are males but is a consideration with larval collections where both sexes are present but not identified.

### Detecting strain-specific assortative mating in field populations

The rationale for using the inbreeding coefficient *F* to assess strain mating behavior is based on the assumption that the frequency of heterozygosity of two SNPs that differ in strain distribution but are otherwise similar (*i.e*., are located in the same exon, have the same neutral polymorphic alternatives, and are detected by the same set of PCR amplification and DNA sequencing reactions) will depend on the frequency of mating between strains relative to that occurring within strains [43].

The frequencies of the C-allele and T-allele for each SNP were estimated using Hardy-Weinberg equilibrium analysis. The allele frequencies for C and T are given by *p* and *q*, respectively, such that *p* + *q* = 1. For each collection *p* was calculated by the equation *p* = freqCC + 0.5[freqY], where freqCC is the observed frequency of CC homozygotes and freqY the observed frequency of Y (CT) heterozygotes. The frequency of the T-allele (*q*) is then given by the equation *q* = 1 - *p*. The local expected heterozygote frequency, *H*_*e*_, is equal to the equation *H*_*e*_ = 2*pq*. The local observed heterozygote frequency, *H*_*o*_, is given by the empirically determined freqY. Wright’s local inbreeding coefficient, *F* = (*H*_*e*_ - *H*_*o*_)/*H*_*e*_, was calculated for SNPs e4_183_ and e4_192_.

The difference in strain-specificity between SNPs e4_183_ and e4_192_ means that heterozygosity at the two sites results from different sets of mating patterns. Heterozygotes at e4_192_ can be produced by mating both between (inter) and within (intra) strains while heterozygotes at the strain-specific e4_183_ site can only occur through interstrain hybridization (Fig. 2A). Therefore, if reproduction barriers between strains exist such that intrastrain is more frequent than interstrain hybridization, then the frequency of observed heterozygosity at e4_183_ will be more severely affected than at e4_192_. This will result in *F*_183_ being greater than *F*_192_.

### DNA sequence and statistical analysis

DNA sequence alignments and comparisons were performed using programs available on the Geneious 10.0.7 software (Biomatters, Auckland, New Zealand). Basic mathematical calculations and generation of graphs were done using Excel and PowerPoint (Microsoft, Redmond, WA). Other statistical analyses including ANOVA, *t*-tests, and *chi*-square were performed using GraphPad Prism version 7.00 for Mac (GraphPad Software, La Jolla California USA).

## Results

Earlier studies demonstrated significant inconsistencies in the distribution of the *COI* and *Tpi* genetic markers in the African fall armyworm populations [9]. A majority of the specimens (all collected from C-strain hosts) displayed the discordant combination of the R-strain *COI*-RS together with the C-strain *Tpi*C marker. This high frequency of discordance between markers was compared to that observed from equivalent Western Hemisphere larval collections also from corn or sorghum hosts (Fig. 3). Specimens from locations in Florida, the Caribbean, and Argentina, each gave similar results to each other, with a mean of 65% showing concordance (*COI*-CS *Tpi*C or *COI*-RS *Tpi*R) in their marker configuration that was significantly higher than the discordant frequency (Fig. 3A). A mean of 4% of specimens was heterozygous for *Tpi*R/*Tpi*C, which is likely to be an underestimate because the larval collections were not identified by gender. Females (ZW) are hemizygous for *Tpi* and so added to the concordant and discordant categories but did not contribute to the number of *Tpi* heterozygotes (Fig. 2B).

**Figure 3.**
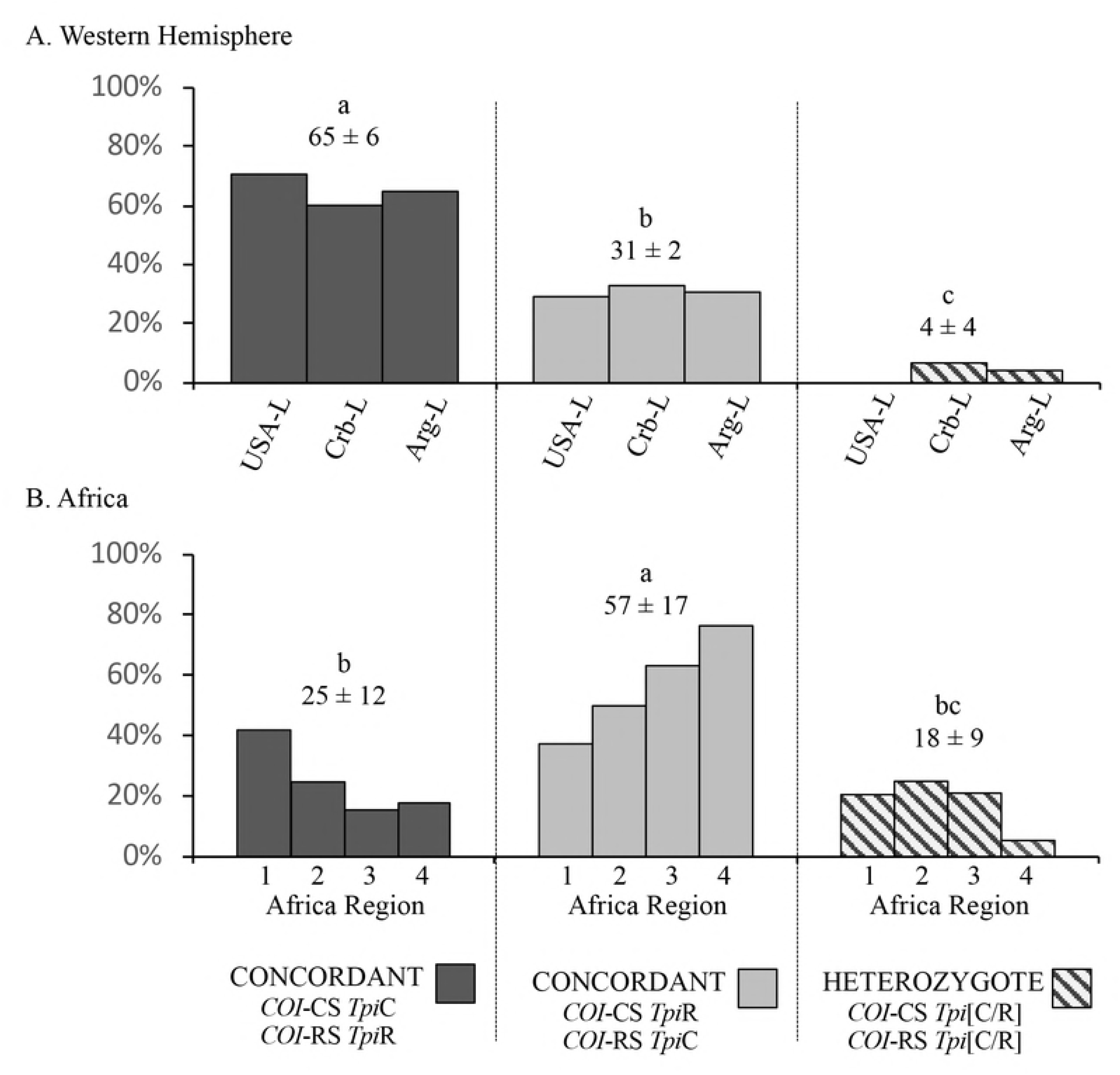
Comparisons of *COI* and *Tpi* marker configurations as categorized by their agreement in identifying strains (concordance), disagreement (discordance), and heterozygosity for the *Tpi* strain-specific alleles (*Tpi*R/*Tpi*C). The mean frequency ± the standard deviation (SD) is listed for each category. Comparisons between the means were done using single-tailed ANOVA analysis (*P* < 0.0001, *R2* = 0.8482, *F* = 16.76) followed by Tukey’s comparison testing with significant differences indicated by different lower-case letters. (A) Western Hemisphere (WH) fall armyworm populations. The categories are composed of the pooled larval collections (Table 1). (B) Africa fall armyworm populations categorized by regions (Fig. 1).

A different pattern was observed with the African fall armyworm populations ([9, 10], Fig. 3B). Only about 25% of the African specimens were concordant for *COI* and *Tpi*, compared to a mean of 57% for the discordant configuration, with both frequencies significantly different from that observed in the Western Hemisphere. The mean of 18% male heterozygotes was 4-fold higher than the mean of 4% observed in the Western Hemisphere. This difference was not statistically significant when analyzed by ANOVA for all three categories (Concordant, Discordant, and Heterozygote), but was significant by a direct pairwise comparison using a two-tailed *t*-test (*P* = 0.0483, *t* = 2.599, *df* = 5).

### Predominance of the *COI*-RS *Tpi*C discordants

A consistent observation from field studies in the Western Hemisphere is that the *COI*-RS *Tpi*C configuration occurs more frequently than the reciprocal discordant genotype of *COI*-CS *Tpi*R [23]. This is illustrated by the Western Hemisphere larval collections from C-strain plants where the proportion of the *COI*-CS *Tpi*R configuration among discordants (27%) was significantly lower than the 73% mean frequency of *COI*-RS *Tpi*C specimens (Fig. 4A). A similar but more extreme trend was found in Africa, where in each of the four regions nearly all discordant specimens were *COI*-RS *Tpi*C (Fig. 4B).

**Figure 4.**
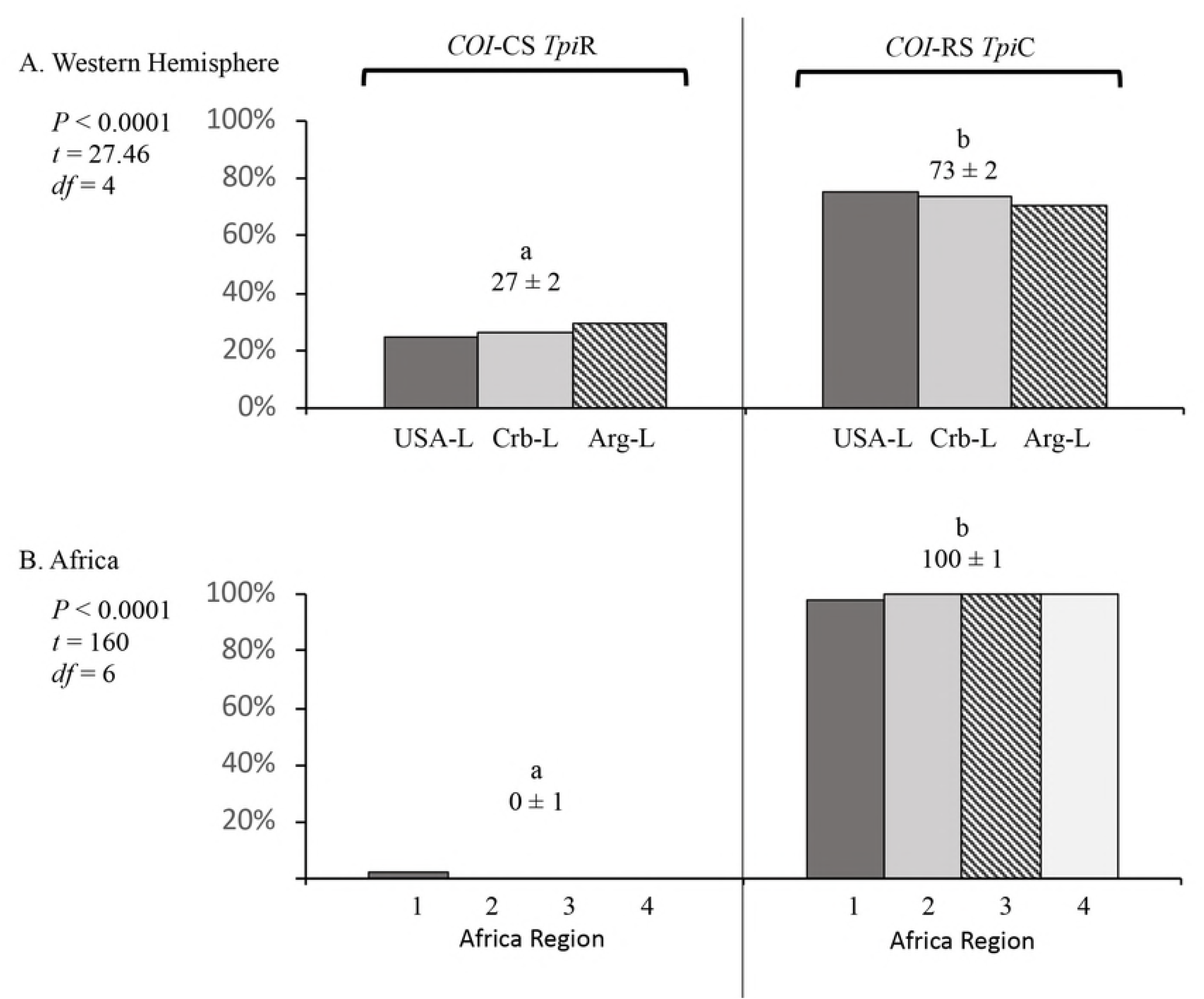
Comparison of the *COI*-CS *Tpi*R and *COI*-RS *Tpi*C configuration frequencies that make up the discordant category described in Fig. 3. The mean frequency ± SD is listed for each category. Pair-wise comparisons between the means were calculated using single-tailed *t*-test with significant differences indicated by different lower-case letters. (A) Western Hemisphere collections. (B) Africa collections. The same collections were used as in Fig. 3. All collections were from corn-strain preferred host plants.

### Detecting strain-specific assortative mating in field populations

Earlier studies demonstrated that comparisons of heterozygosity frequencies of SNPs that differ in their degree of strain-specificity could provide evidence of reduced interstrain hybridization in field populations [43]. The initial studies were based on pheromone collections from the same location and time period, which were presumed to represent a single mating population with the allele frequencies observed used to calculate the inbreeding coefficient, *F*. Such collections consistently showed a higher *F* value for the strain-specific e4_183_ SNP (*F*_183_) compared to that of the non-specific e4_192_ SNP (*F*_192_) independent of location (Fig. 5A).

**Figure 5.**
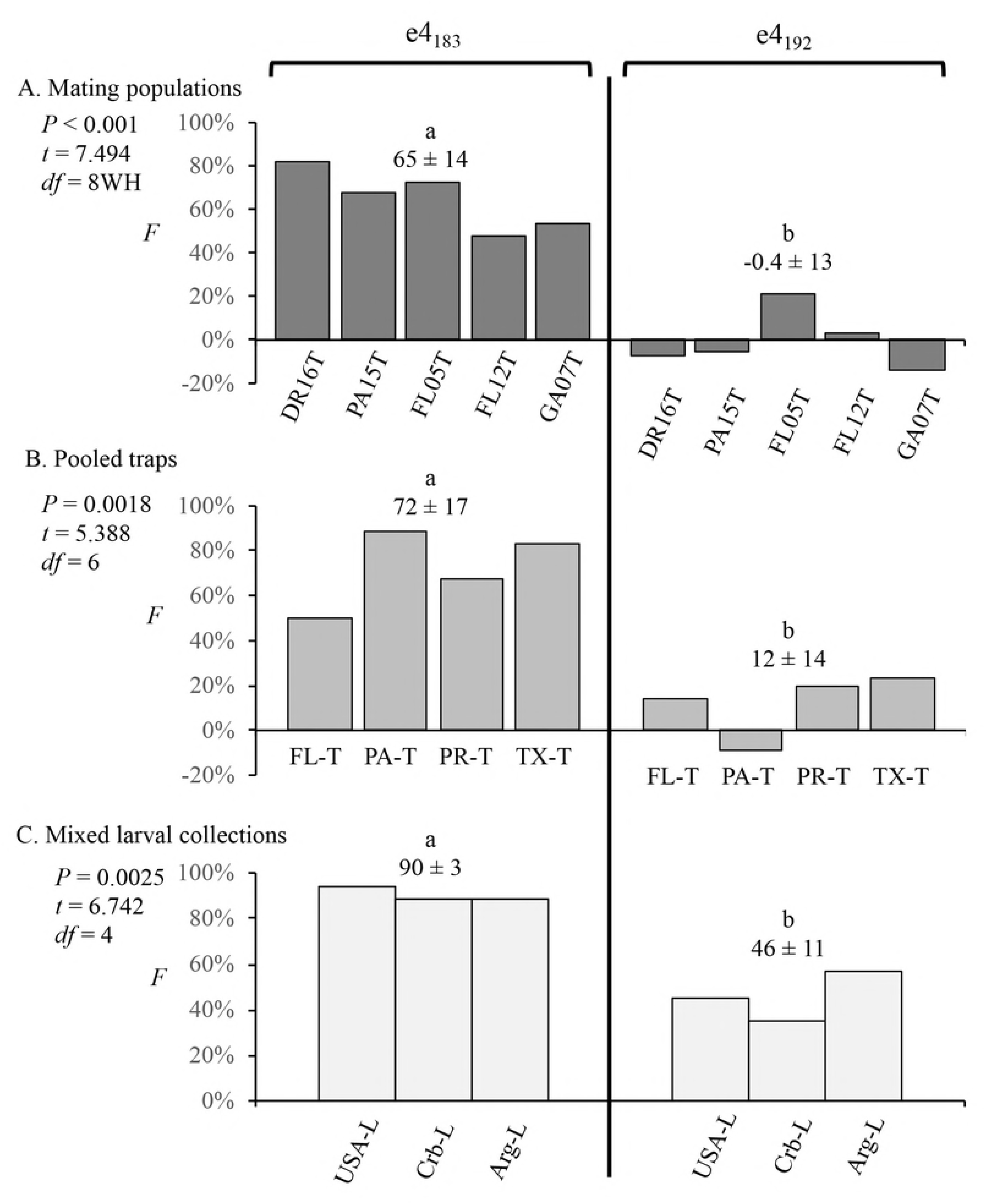
Comparisons of inbreeding coefficients (*F*) for the e4_183_ and e4_192_ SNPs calculated for different mating populations and collection methods. The mean *F* ± SD is listed for each category. Pairwise comparisons between the means were calculated using single-tailed t-test with significant differences indicated by different lower-case letters. (A) Inbreeding coefficients calculated for pheromone trap collections from a single time and location and therefore are representative of a single breeding population. (B) Inbreeding coefficients calculated for pooled pheromone trap collections from different times or locations within a state. (C) Inbreeding coefficients calculated for pooled larval collections from a given country or region.

A characteristic of this technique, at least in principle, is that external factors can be discounted as long as they equally affect both *F* values such that their relative difference is maintained. This was confirmed in two ways. In the first, pheromone trap collections from different times and locations were pooled resulting in allele frequencies derived from multiple mating populations (Fig. 5B). In this case the calculated composite *F* coefficients were not representative of an actual field population but still consistently showed *F*_*183*_ greater than *F*_*192*_ at all locations. A second test pooled larval collections from different time periods. Because the gender of the larvae was unknown, an unspecified subset of these will be female, which have only one copy of the *Tpi* gene and are indistinguishable from males that are homozygous for a *Tpi* allele. Since *Tpi* heterozygotes are only possible in males, the inclusion of female larvae will result in an underestimate of heterozygote frequency and therefore an overestimate of *F*. The results of the larval collections were consistent with these expectations, with both *F*_183_ and *F*_192_ values generally higher than observed with the trap collections, but with *F*_*183*_ still significantly greater than *F*_*192*_ (Fig. 5C).

This methodology was applied to two different sets of African collections. The first were pheromone trap collections from four different time periods at a single location in Togo, with each time period collection assumed to be representative of the local mating population. There was substantial variation in *F* values between sites but no significant difference between mean *F*_183_ and *F*_192_ (Fig. 6A). Similar results were obtained with the second collection set that was composed of pooled larval specimens from multiple locations within each of the four African regions (Fig. 6B). Once again there was no statistically significant difference between *F*_183_ and *F*_192_ values. The African larval collections also differed from its Western Hemisphere counterpart in that the expected overestimate of *F* (in larval collections that presumably includes females) was not observed (compare Fig. 5C and 6B). The reason for this difference is unknown.

**Figure 6.**
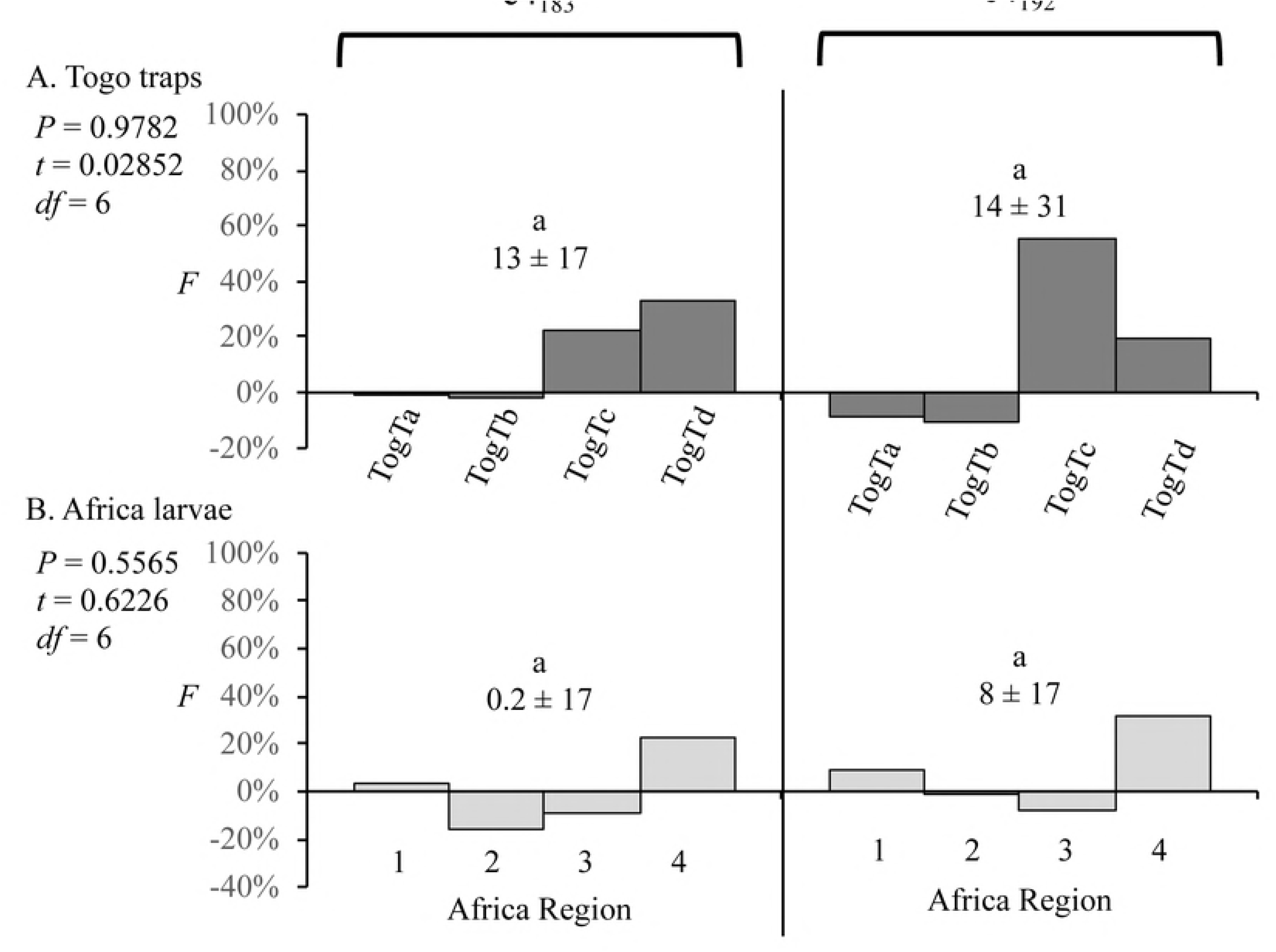
Comparisons of inbreeding coefficients (*F*) for the e4_183_ and e4_192_ SNPs in African fall armyworm collections. The mean *F* ± SD is listed for each category. Comparisons between the means were done using single-tailed t-test with significant differences indicated by different lower-case letters. Comparisons between the means were done using single-tailed t-test and no significant differences were found. (A) Inbreeding coefficients calculated for pheromone trap collections Togo. (B) Inbreeding coefficients calculated for pooled larval collections from four different regions in Africa (Fig. 1).

## Discussion

The inbreeding coefficient *F* describes the difference between the observed heterozygosity (*H*_*o*_) of a particular allele and the heterozygosity expected (*H*_*e*_) as calculated from the parental allele frequencies and based on assumptions of random mating and equal fitness of the resultant products. In the absence of reproductive barriers between parents the expected relationships will be *H*_*o*_ = *H*_*e*_ and *F* = 0, while the existence of reproductive barriers will result in *H*_*o*_ < *H*_*e*_ and *F* approaching +1. This methodology was used in a previous study to demonstrate the existence of reproductive barriers between fall armyworm host strains that could be detected in field populations. It showed that *F* for strain-specific e4_183_ SNP on average 4-fold larger than the *F* of the nearby e4_192_ SNP that is less strain-specific, consistent with a significantly reduced frequency of interstrain hybridization [43].

In this study the method was shown to be viable in composite collections from different time periods and locations, even when using larval specimens of undetermined gender. This allows for the pooling of collections with small sample sizes, making possible analysis of the larval African specimens grouped by geographical region. The one limitation are collections where only a single parental allele is present as heterozygotes in these cases are not possible.

### Are both fall armyworm strains present in Africa?

The presence of the R-strain in Africa is based on mitochondrial markers primarily in the *COI* gene [8-11], but is made uncertain by three observations. First, there are widespread reports throughout the sub-Sahara region of fall armyworm infestation in corn and sorghum, two hosts preferred by the C-strain, but major infestations in host plants preferred by the R-strain have yet to be documented. Second, over 95% of the specimens tested from multiple locations in Africa expressed the C-strain diagnostic *Tpi*C marker [9, 10], indicating that the R-strain population as defined by *Tpi* is rare. Third, and more problematic, is that the majority of the *Tpi*C specimens express the R-strain *COI*-RS haplotype. These observations indicate that the *COI* strain markers are in substantial disagreement with both host plants and *Tpi*. Recently, two other mitochondrial genes (*cyt b* and *COIII*) were characterized in fall armyworm collected from Uganda and shown to exhibit strain-specific SNPs [11]. The strain identity defined by the *cyt b* and *COIII* markers were consistent with each other and with those found in *COI*. The equivalence of these mitochondrial markers with respect to identifying host strains indicates that the disagreement with *Tpi* goes beyond *COI* and involves multiple segments of the mitochondrial genome.

If the mitochondrial haplotypes are not reliable markers of strain identity in the African fall armyworm populations then there is little current evidence supporting the presence of an African R-strain. The absence of a significant R-strain population would be consistent with the comparative heterozygosity *F* study that found no evidence of strain-specific mating barriers in the Africa populations. In summary, at this point the best available evidence is that the R-strain is not present in significant numbers in Africa. The question then is what happened to the *COI* marker?

One possibility is that the African fall armyworm population is made up of the C-strain and interstrain hybrids, with the R-strain either absent or too low to be detected by the comparative heterozygosity studies. The rationale for this proposal is outlined in Figure 7. Discordance between *COI* and *Tpi* is presumed to have occurred due to interstrain crosses, which can occur in two directions with each producing the *Tpi*R/*Tpi*C heterozygote (Fig. 7A, B). Subsequent mating of the hybrids to each other or a backcross to the parental male will generate the discordant configurations. Figure 7C shows the Africa haplotype data (from Fig. 3) recalculated to estimate the proportions of only the male specimens, which should provide a more accurate assessment of heterozygote frequency. This was done by assuming a 1:1 sex ratio and reducing the concordant and discordant numbers by 50%. The results show that 28% of the African fall armyworm are interstrain heterozygotes, which is more than the proportion of *COI*-CS *Tpi*C C-strain (20%). The predominant genotype is the *COI*-RS *Tpi*C discordant that is present at 51%. Therefore, in this scenario the large majority (about 80%) of fall armyworm in Africa are derived from an interstrain hybridization event sometime in their past, with “true” C-strain (*COI*-CS *Tpi*C) a minority and the “true” R-strain (*COI*-RS *Tpi*R) mostly absent.

**Figure 7.**
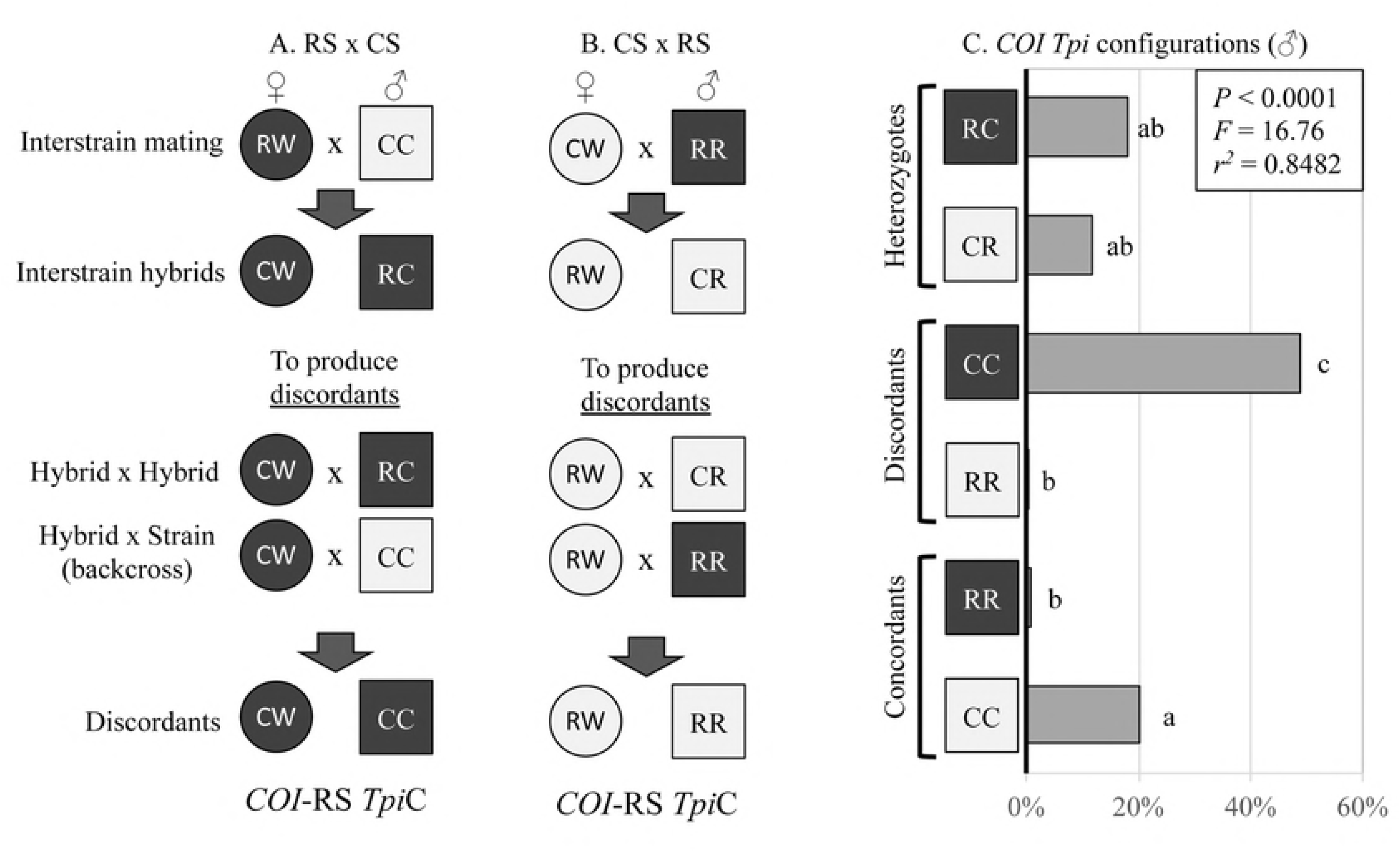
Mating scenarios to produce *Tpi* heterozygotes and *COI Tpi* discordant genotypes their relative frequencies in extrapolated male specimens from Africa. The *Tpi*R (R) and *Tpi*C (C) alleles are defined by the e4_183_ polymorphism. Females are designated by a circle and are hemizygous for *Tpi* and so can only carry one *Tpi* allele, while males (square) and are either homozygous (RR, CC) or heterozygous (RC, CR) for *Tpi*R and *Tpi*C. Dark shading indicates a *COI*-RS mitochondrial haplotype while light shading identifies *COI*-CS. A) Female rice-strain (RS) mating male corn-strain (CS), while B) represents the reciprocal cross. C) Graph of the frequencies of each *COI Tpi* configuration within the male subset of the data described in Fig. 3B (see text for explanation). Lower case letters indicate significant differences as determine by ANOVA and Tukey’s multiple comparison test.

### Can e4_183_ heterozygotes be assumed to be interstrain hybrids?

A major assumption in this proposal is that heterozygosity at e4_183_ is an accurate marker for interstrain hybrids, which genetically is defined by heterozygosity at all the loci critical for driving strain divergence. There are two lines of evidence in support. The first and most compelling is that e4_183_ is an accurate marker of strain identity as indicated by surveys at multiple locations and times in the Western Hemisphere [18, 23, 44, 45, 50, 51]. This indicates a strong linkage of this SNP to the genes most critical for strain differentiation, directly implicating functions on the *Z*-chromosome located near *Tpi* by recombination distance. The second is the strain-specificity exhibited by mitochondrial markers that has been confirmed by multiple studies in the Western Hemisphere [5, 6, 19, 52]. Assuming the mitochondria are not involved in strain differentiation, this suggests a coincidental association that links the maternal inheritance pattern of mitochondria with the sex-specific inheritance of the *Z* and *W* chromosomes. In other words, the strain specificity of the *COI* and *Tpi* markers indicate that autosomal genes play at best a secondary role in strain differentiation. Since most sex-linked functions are located on the *Z*-chromosome rather than *W*, a *Z*-linked marker like e4_183_ should act as an accurate proxy for true interstrain hybrids.

### Why do the African and Western Hemisphere populations differ?

Descriptions of how fall armyworm was introduced into Africa are completely speculative at this time but there are plausible scenarios for how an interstrain hybrid dominated population could have emerged. In the Western Hemisphere the distribution of the two strains is largely sympatric, particular in the habitats associated with C-strain preferred host plants [5, 6, 19, 52]. It is therefore likely that a contaminated shipment would carry both strains. Such a small invasive propagule would be subject to the variety of effects associated with population bottlenecks, most notably inbreeding depression due to recessive deleterious alleles. This will tend to reduce genetic variation and fitness within the normal breeding groups, which in this case are the two strains, thereby tending to increase the proportion of progeny produced by interstrain crosses. If this promotion of hybrid frequencies is greater than the reproductive barriers that limit productive mating between strains, it is plausible that a hybrid population could emerge and predominate.

The expected proportions of the two strains would depend on multiple factors that include the initial composition of the invasive propagule, the availability of host plants, and subsequent stochastic events. If the initial contaminated product was a C-strain host, a majority C-strain in the initial introductory population is likely, increasing the probability of its survival in the established population and conversely the marginalization and eventual loss of the R-strain.

### What characteristics are expected in the hybrid and discordant genotypes?

The behavior of interstrain hybrids is unknown. The heterozygotes so far captured in Africa have all been from corn or sorghum indicating attraction to C-strain hosts, while the absence of reproductive barriers as assayed by the comparative heterozygosity analysis suggests an unrestricted capacity to mate with *Tpi*R moths.

It is noteworthy that the *COI*-RS *Tpi*C class is much greater than the reciprocal *COI*-CS *Tpi*R discordants in Africa, just as it is in the Western Hemisphere. This indicates that the set of matings initiated by female R-strain crossed to male C-strain (Fig. 7A) is much more frequent and/or productive than the reciprocal (Fig. 7B). This observation does not appear to be consistent with other findings indicating that the female R-strain crossed to male C-strain mating (but not the reciprocal) produced daughters with significantly reduced fertility [39]. The reason for this apparent inconsistency is unknown but suggests the possibility of unknown complexity and variability in the nature of the reproductive barriers between strains.

## Conclusions

Consistent findings of discrepancies between the *COI* and *Tpi* strain markers and the absence of strain-specific mating barriers indicate that the R-strain subpopulation found in the Western Hemisphere is currently not present in significant numbers in Africa. This could have important consequences in assessing the threat of fall armyworm to different crop species. Evidence is presented that suggest the Africa fall armyworm populations is primarily composed of interstrain hybrids with a C-strain minority population.

## Acknowledgements

Thank you to Dr. Jean Thomas (USDA-ARS) and Dr. Robert Meagher (USDA-ARS) for advice on the manuscript. The use of trade, firm, or corporation names in this publication is for the information and convenience of the reader. Such use does not constitute an official endorsement or approval by the United States Department of Agriculture or the Agricultural Research Service of any product or service to the exclusion of others that may be suitable.

